# Tutorial: Investigating SARS-CoV-2 evolution and phylogeny using MNHN-Tree-Tools

**DOI:** 10.1101/2021.12.21.473702

**Authors:** Thomas Haschka

## Abstract

The Covid-19 pandemic has caused at more than 3 million deaths by Mai this year [1]. It had a significant impact on the daily life and the global economy [2]. The virus has since its first recorded outbreak in China [3] mutated into new strains [4]. The Nextstrain [5] project has so far been monitoring the evolution of the virus. At the same time we were developing in our lab the MNHN-Tree-Tools [6] toolkit, primarily for the investigation of DNA repeat sequences. We have further extended MNHN-Tree-Tools [6] to guide phylogenetics. As such the toolkit has evolved into a high performance code, allowing for a fast investigation of millions of sequences. Given the context of the pandemic it became evident that we will use our versatile tool to investigate the evolution of SARS-CoV-2 sequences. Our efforts have cumulated in this tutorial that we share with the scientific community.

## 1 Introduction

The SARS-CoV-2 virus was first recorded in China [3] in 2019. At least 3 million deaths caused by the virus [1] have been recorded by Mai 2021. The virus further has impacted social life and economies around the globe [2]. Even though successful vaccine campaigns are run, especially in the western world, the virus has since underwent several escape mutations that might jeopardize them in the future [4]. As such constant monitoring of the viruses evolution is needed. As the virus sequence databases are growing faster, more efficient solutions are needed in order to rapidly infer phylogeny. More than 1.5 million sequenced samples are openly available to researchers and can be readily downloaded from the internet. Having our in house developed MNHN-Tree-Tools [6] toolkit at hand, we grasped this opportunity and investigated the diversity of the virus. In order to yield proper results, the sequences that we prepare for our algorithm should at least roughly correspond to the sequence distribution that is spread around the globe. So far we have no means to verify if this might be the actual case. The data that is freely and openly available to us overrepresents SARS-CoV-2 sequences for the United States. Aware of these shortcomings we proceeded in our endeavor and found that we can clearly distinguish variants of concern, and that clades, especially pointing to the alpha and delta variants, are clearly visible in tree traced by the MNHN-Tree-Tools algorithm. Such a tree is outlined in figure 1. We like to point out that our trees monitor the evolution of at least 29 500 bases pairs of the SARS-CoV-2 virus and do not limit themselfs to regions that express the spike protein. We further find that according to our dataset alpha, beta and gamma variants seem to be closer related and that the delta variant seems to envolve apart. These properties are visibly in the trees that we have traced shown in detail in figures 1, 2 and 3.

**Figure 1:**
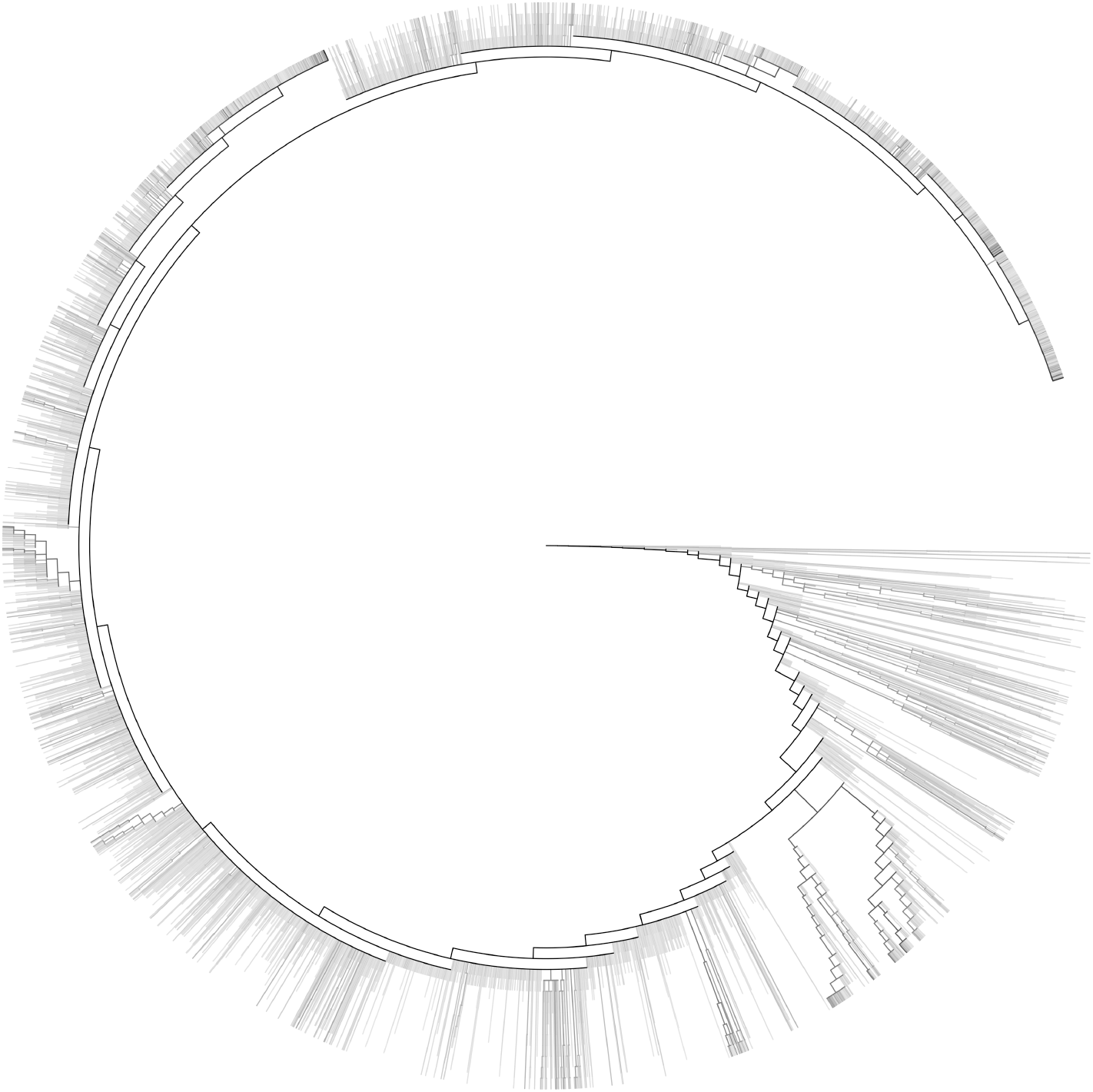
The complete MNHN-Tree-Tools infered tree of our downloaded dataset of 1 442 669 SARS-CoV-2 sequences. Number of sequences in nodes are shown logarithmically in a gradient from with (0 sequences) to dark black (1 442 669).

**Figure 2:**
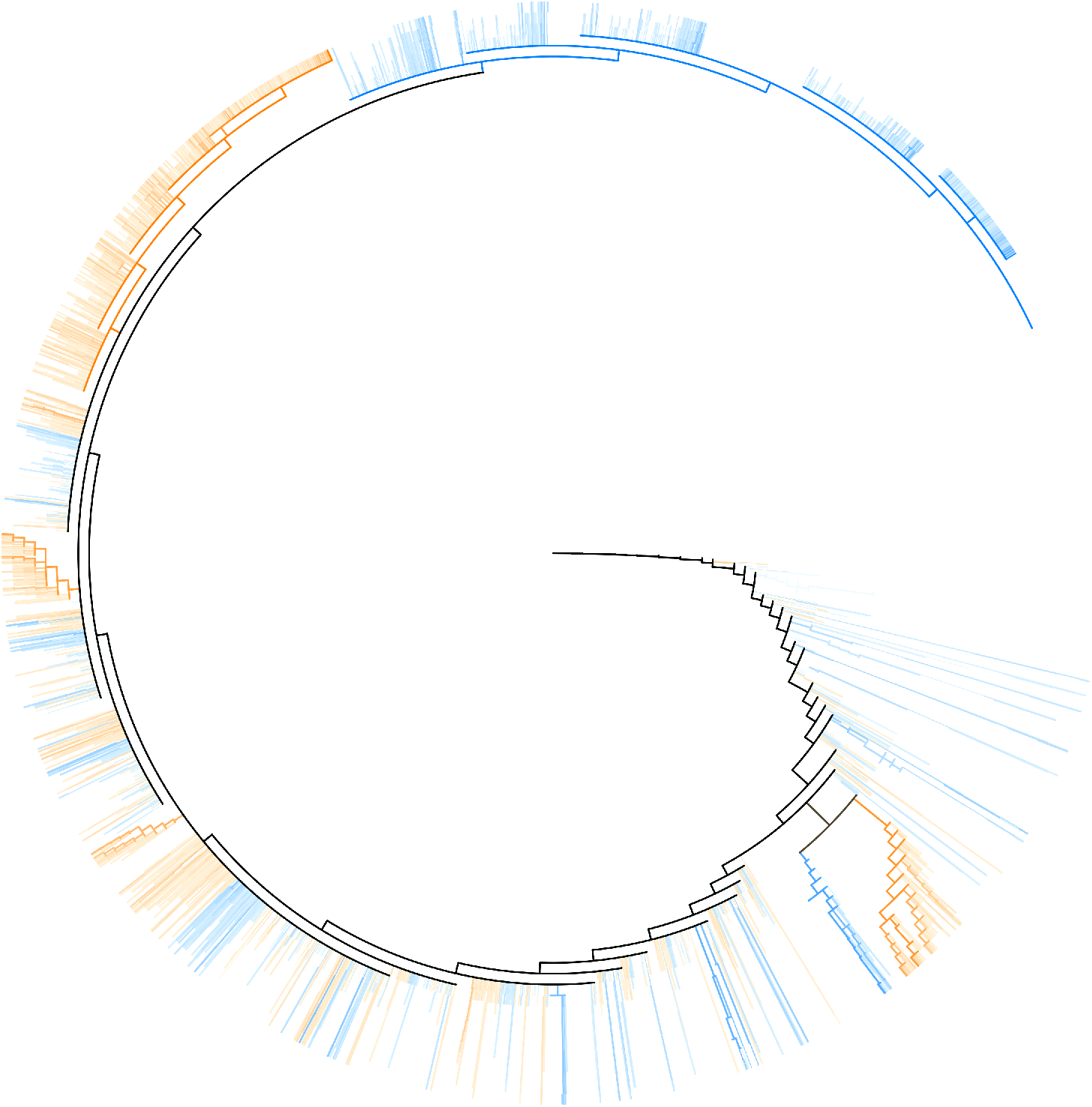
A tree composed of only alpha and delta variant sequences. Delta variant sequences are highlighted in orange, Alpha sequences are highlighted in blue. Mixed clusters are outlined as gray-black. The number of sequences is highlighted by a logarithmic color intensity gradient from white clusters containing no sequences to increasing saturation up until all sequences are contained in full black.

**Figure 3:**
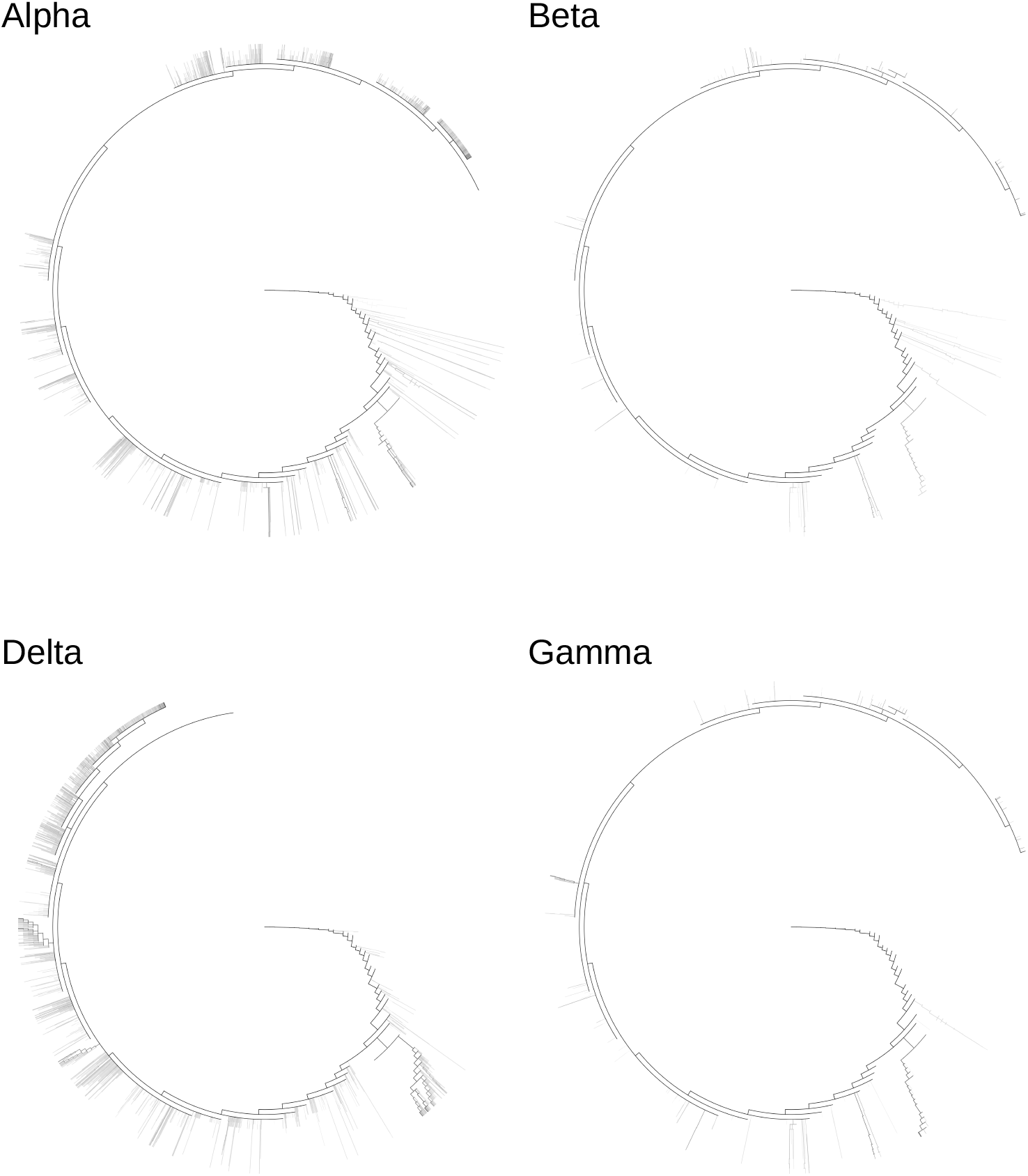
Partial highlighted parts of the tree outlined in figure 1. Highlighted are for the variants alpha, beta, gamma and delta the nodes that contain sequences of the corresponding variant. The nodes highlighte the number of sequences of the variant in questions in logarithmic gradient from white for no occurrence of the variant to black, all sequences found in the dataset and attributed to the variant in question are part of this node.

## 2 Tutorial - Methods

### 2.1 Machine and software requirements

This tutorial was performed on a 64bit Intel (R) architecture Linux machine consisting of 28 real and due to hyper threading 56 virtual cores. Due to the nature MNHN-Tree-Tools it is imperative that the CPU cores of the machine support the AVX instruction set. The machine further was equipped with 128 gigabytes of random access memory (RAM). MNHN-Tree-Tools was installed on the machine as described in its manual^1^ [6]. We suggest that the reader following this tutorial uses a similar machine, although different configurations might work.

### 2.2 Covid-19 sequence retrieval

On the 16th of september 2021 we downloaded 1 442 669 nucleotide sequences of the SARS-CoV-2 virus from the NCBI SARS-CoV-2 Resources website ^2^. The sequences were selected to match a minimum length of 29 500 bases pairs and do not exceed a length of 32 300 bases pairs. Further only sequences obtained from human, homo sapiens, host identified by taxid:9606 in the database were selected. We instructed the NCBI website to prepare the data in FASTA format, containing only the SARS-CoV-2 Pango linage identifier as well as the NCBI accession code in the sequence identifier lines of the FASTA file. No further filtering of the FASTA raw resources was performed. The obtained FASTA file is stored as *covid-ncbi-16-sept-2021.fasta*.

### 2.3 K-merization of the Covid-19 sequences

Having these 1 442 669 nucleotide sequences at hand we went ahead and instructed MNHN-Tree-Tools to calculate the k-mer represenations of these sequences. In order to capture the information held by sequences of a length of approximately 32 000 bases pairs we opted for a 6-mer representation. The transformation was performed using the *fasta2kmer* tool by executing the following command:

~~~
fasta2kmer covid -ncbi -16 - sept -2021. fasta 6 56 0 > covid -ncbi -16 - sept -2021.6 mer
~~~

which transforms all downloaded SARS-CoV-2 sequences stored in *covid-ncbi-16-sept-2021.fasta* to 6mer sequences, using 56 compute threads.

### 2.4 Projections onto Principal Components

In the next step we used the *kmers2pca* tool of the MNHN-Tree-Tools toolbox to perform PCA on the 6-mer representations of the SARS-CoV-2 sequences. The first 7 principal components with the 7 highest corresponding eigenvalues were selected. Even though information capture at this reduced number of principal components might not be optimal we stopped at this resolution in order to avoid the course of dimensionality that might hinder subsequent clustering runs. All 6-mer representations were projected into the 7 dimensions spun by the selected 7 principal components. The *kmer2pca* tool from MNHN-Tree-Tools was used, as outlined below, in order to perform this task:

~~~
kmer2pca covid -ncbi -16 - sept -2021.6 mer covid -ncbi -16 - sept -2021.6 mer - pca7v \
covid -ncbi -16 - sept -2021.6 mer -ev 7 56
~~~

### 2.5 Adaptive clustering and tree building

Using the *adaptive_clustering_PCA* tool part of MNHN-Tree-Tools we continue performing clustering using the DBSCAN [7] algorithm at different sequence densities and subsequent treebuilding. For in depth details we refer the reader to the algorithmic supplement published with the article [6]. The parameters chosen for the run were: Initial epsilon *ϵ*_0_ = 0.5, an epsilon increase per step Δ*ϵ* = 0.1 and a minimal number of sequences to form or extend a cluster of minpts = 6, which resulted in a tree containg 50 stages. The following commands were executed to perform this task:

~~~
mkdir clusters
adaptive_clustering_PCA covid -ncbi -16 - sept -2021. fasta 0.5 0.1 6 \
clusters /covid -ncbi -16 - sept -2021 -6 mer -pca -7v -0.5 -0.1 -6 - 56 7 \
covid -ncbi -16 - sept -2021.6 mer - pca7v > \
clusters /covid -ncbi -16 - sept -2021 -6 mer -pca -7v -0.5 -0.1 -6 - output
~~~

In order to visualize the results we have to create a tree in Newick format^3^ from the adaptive clustering run that we have performed beforehand. This is done by invoking *split_sets_to_newick* as follows:

~~~
mkdir clusters /covid -ncbi -16 - sept -2021 -6 mer -pca -7v -0.5 -0.1 -6 - additions
cd clusters
split_sets_to_newick 0 0 \
covid -ncbi -16 - sept -2021 -6 mer -pca -7v -0.5 -0.1 -6 -00* > \
covid -ncbi -16 - sept -2021 -6 mer -pca -7v -0.5 -0.1 -6 - additions /full - tree . dnd
~~~

The resulting *full-tree.dnd* file allows us to use common software tools that visualize Newick trees such as the Newick utilities [8]. The resulting tree is shown in its entirety in figure 1. The different shades are proportional to the logarithm of the number of sequences found in the tree nodes. From white for 0 sequences, to full black for the 1 442 669 sequences contained in our dataset. In order to visualize the tree with the shading outlined above a *map* that can be supplied to the Newick utilities [8] has to be created. The *tree_map_for_split_set* tool using the final tree layer, containing all sequences, in our case number 49, is perfect for such a task:

~~~
tree_map_for_split_set covid -ncbi -16 - sept -2021 -6 mer -pca -7v -0.5 -0.1 -6 -0049 \
../ covid -ncbi -16 - sept -2021. fasta 1 2 1 ,#000000 \
covid -ncbi -16 - sept -2021 -6 mer -pca -7v -0.5 -0.1 -6 -00* > \
covid -ncbi -16 - sept -2021 -6 mer -pca -7v -0.5 -0.1 -6 - additions /full -log - gradient . map
~~~

### 2.6 Visulization of the different Covid-19 variants

In the dataset lineages B.1.1.7 Q.1 Q.2 Q.3 Q.4 Q.5 Q.6 Q.7 and Q.8 were designated to be of the alpha, B.1.135 of beta, P.1 to be of gamma and B.1.617.2 AY.1 AY.2 AY.3 AY.3.1 AY.4 AY.4.1 AY.5 AY.5.1 AY.5.2 AY.6 AY.7 AY.7.1 AY.7.2 AY.8 AY.9 AY.10 AY.11 AY.12 AY.13 AY.14 AY.15 AY.16 AY.17 AY.17 AY.19 AY.20 AY.21 AY.22 AY.23 AY.24 AY.25 AY.26 AY.27 AY.28 AY.29 AY.31 AY.32 AY.33 AY.34 AY.35 AY.36 AY.37 and AY.38 of the delta variant. Using the FASTA file annotations for the *split_set_from_annotation* tool which is part of MNHN-Tree-Tools were created. We first processed the downloaded FASTA file with general shell utilities available on our Linux system. In order for the following commands to work the reader has to make sure that Pango linage is in the “$1” row according to the *awk* tool in the FASTA annotation lines.

~~~
grep ’>’ covid -ncbi -16 - sept -2021. fasta > covid -ncbi -16 - sept -2021. annotations
awk ’{print $1}’ covid -ncbi -16 - sept -2021. annotations > \
covid -ncbi -16 - sept -2021. strains
~~~

The commands above extract all Pango lineage codes into a text file. In order to create valid annotation from this text file we created a small utility in C whose source code is outlined in the following:

~~~
# include < stdio .h>
# include < string .h>
int main (int argc , char ** argv ) {
  int i;
  int check ;
  FILE * f = fopen (argv [1] ,“r ”);
  int n_words = argc -2;
  char string_buffer [200];
  while (1 == fscanf (f ,“% s”, string_buffer )) {
   check = 0;
   if (strlen (string_buffer) > 1) {
    for (i =0;i< n_words ;i ++) {
     if (0 == strcmp ((argv [i +2]) ,(string_buffer +1))) {
      check = 1;
     }
    }
   }
   printf ("% i\n", check );
 }
 return (0);
}
~~~

This tool allows us to create annotation files containing either 1 if a certain sequence belongs to a certain variant, or 0 if it does not. We invoke our utility like the following to create annotations for the alpha, beta, gamma and delta variants:

~~~
./a. out covid -ncbi -16 - sept -2021. strains \
B .1.1.7 Q.1 Q.2 Q.3 Q.4 Q.5 Q.6 Q.7 Q.8 > covid -ncbi -16 - sept -2021. alpha
./a. out covid -ncbi -16 - sept -2021. strains B .1.351 > covid -ncbi -16 - sept -2021. beta
./a. out covid -ncbi -16 - sept -2021. strains P.1 > covid -ncbi -16 - sept -2021. gamma
./a. out covid -ncbi -16 - sept -2021. strains B .1.617.2 AY .1 AY .2 AY .3 \
  AY .3.1 AY .4 AY .4.1 AY .5 AY .5.1 AY .5.2 AY .6 AY .7 AY .7.1 AY .7.2 AY .8 \
  AY .9 AY .10 AY .11 AY .12 AY .13 AY .14 AY .15 AY .16 AY .17 AY .17 AY .19 \
  AY .20 AY .21 AY .22 AY .23 AY .24 AY .25 AY .26 AY .27 AY .28 AY .29 AY .31 \
  AY .32 AY .33 AY .34 AY .35 AY .36 AY .37 AY .38 > \
  covid -ncbi -16 - sept -2021. delta
~~~

Using the sets created with this tool that indexed the sequences by the afformentioned annotations for alpha, beta, gamma and delta variants we are able to use the *tree_map_split_set* tool, again part of MNHN-Tree-Tools, to create color maps that allow us to highlight the different variants in our tree using the newick utilities [8]. We performed this step using the following command on the BASH shell:

~~~
for i in alpha beta gamma delta
do tree_map_for_split_set ../ covid -ncbi -16 - sept -2021. $i -split -set \
../ covid -ncbi -16 - sept -2021. fasta 1 2 2 ,#000000 \
covid -ncbi -16 - sept -2021 -6 mer -pca -7v -0.5 -0.1 -6 -00* > \
covid -ncbi -16 - sept -2021 -6 mer -pca -7v -0.5 -0.1 -6 - additions /$i - opposition - map
done
~~~

All trees were finally visualized using the newick utilities [8]. A multicolored tree highlighting both the alpha and gamma variant in a single tree was created by overlaying and postprocessing the results using the GIMP program. The resulting visualizations are highlighted in figures 1,2 and 3.

## 3 Discussion

The data that we obtained is probably skewed as most of the sequences are either of the alpha or delta variant, and most sequences in the dataset originate from the United States with way more populus countries such as India significantly underrepresented. Intrigingly we find very well that beta and gamma variants are mostly related to clades in which the alpha variant labeled sequences reside. Looking at figures 1, 2 and 3 we clearly can identify on the upper left a delta clade and on the upper right an alpha clade. This finding further is replicated by two clades on the lower right side of the figures 1, 2 and 3. In this lower right area the tree develops two boot like structures, where the left boot contains alpha, but intriguingly also beta and gamma sequences, while the right boot contains only delta sequences. A tree provided by the https://nextstrain.org website, shown in figure 4, looks different, but seems to highlight the same features in the sense that alpha, beta and gamma sequences seem closer related than delta sequences. An in depth comparison between our algorithm, tree and data with the one provided by Nextstrain [5], albeit interesting lies out of the scope of this tutorial. The interested reader is referred to the algorithmic supplement provided with MNHN-Tree-Tools [6].

**Figure 4:**
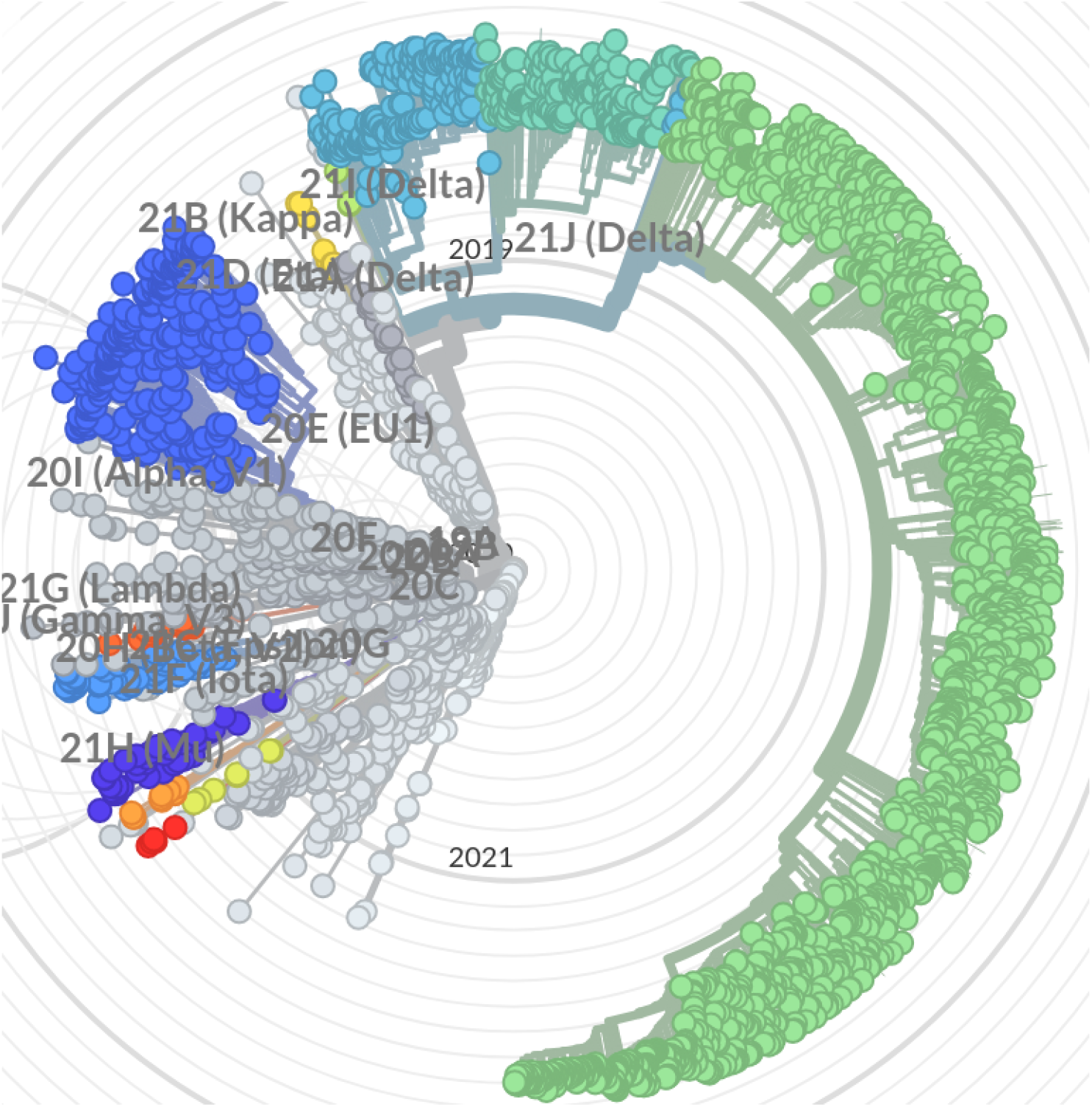
Clades of SARS-CoV-2 strains as infered from openly available data on the https://nextrain.org website.

We conclude that MNHN-Tree-Tools and its efficiency handling millions of sequences can be useful in studying the evolution of SARS-CoV-2 and its phylogenetics. The public dataset available here provides fast results. These results should however be clearly inspected, and readers are referred to use more comprehensive and cleaned data with this algorithm.

## 4 Acknowledgments

We thank the french Musém National d’Histoire Naturelle (MNHN) for providing the computational resources in order to write this tutorial.

## Footnotes

1 http://treetools.haschka.net/

2 https://www.ncbi.nlm.nih.gov/sars-cov-2/

3 https://evolution.genetics.washington.edu/phylip/newicktree.html

## References

[1] Owen Dyer. Covid-19: Study claims real global deaths are twice official figures. BMJ, 373, 2021.

[2] Samuel Asumadu Sarkodie and Phebe Asantewaa Owusu. Global assessment of environment, health and economic impact of the novel coronavirus (covid-19). Environment, Development and Sustainability, 23(4):5005–5015, Apr 2021.

[3] Kathy Leung, Joseph T Wu, Di Liu, and Gabriel M Leung. First-wave covid-19 transmissibility and severity in china outside hubei after control measures, and second-wave scenario planning: a modelling impact assessment. The Lancet, 395(10233):1382–1393, 2020.

[4] Md Kamal Hossain, Majid Hassanzadeganroudsari, and Vasso Apostolopoulos. The emergence of new strains of sars-cov-2. what does it mean for covid-19 vaccines? Expert Review of Vaccines, 20(6):635–638, 2021. PMID: 33896316.

[5] James Hadfield, Colin Megill, Sidney M Bell, John Huddleston, Barney Potter, Charlton Callender, Pavel Sagulenko, Trevor Bedford, and Richard A Neher. Nextstrain: real-time tracking of pathogen evolution. Bioinformatics, 34(23):4121–4123, 05 2018.

[6] Thomas Haschka, Loic Ponger, Christophe Escudé, and Julien Mozziconacci. MNHN-Tree-Tools: a toolbox for tree inference using multi-scale clustering of a set of sequences. Bioinformatics, 37(21):3947–3949, 06 2021.

[7] Martin Ester, Hans-Peter Kriegel, Jörg Sander, and Xiaowei Xu. A density-based algorithm for discovering clusters a density-based algorithm for discovering clusters in large spatial databases with noise. In Proceedings of the Second International Conference on Knowledge Discovery and Data Mining, KDD’96, page 226–231. AAAI Press, 1996.

[8] Thomas Junier and Evgeny M. Zdobnov. The Newick utilities: high-throughput phylogenetic tree processing in the Unix shell. Bioinformatics, 26(13):1669–1670, 05 2010.

